# Investigation on the effect of sonic stimulation on *Xanthomonas campestris* at the whole transcriptome level

**DOI:** 10.1101/607663

**Authors:** Pooja Patel, Chinmayi Joshi, Vijay Kothari

**Affiliations:** Institute of Science, Nirma University, Ahmedabad- 382481, India

**Keywords:** Transcriptome, Xanthan, Exopolysaccharide (EPS), Sonic stimulation

## Abstract

A gram-negative bacterium *Xanthomonas campestris* was subjected to sonic stimulation with sound pertaining to 1000 Hz at three different sound intensities. The *X. campestris* culture subjected to sonic stimulation at 66 dB produced 1.69 fold higher exopolysaccharide. Whole transcriptome analysis of this sonic-stimulated culture revealed a total of 115 genes expressed differentially in the sonic-stimulated culture, majority of which were coding for different proteins including enzyme. This study demonstrates the property of the test bacterium of being responsive to sonic/vibrational stimulation.

## 1. Introduction

Our knowledge regarding whether and how microorganisms respond to sonic frequencies (i.e. sound pertaining to human audibility) is very limited. Hitherto this has largely remained an under-investigated area. We and few other researchers in past have indicated that microbes can get affected by sonic stimulation [Ying et al., 2009; Shaobin et al., 2010; Aggio et al., 2012; Kim et al., 2016; Liu et al., 2016; Murphy et al., 2016; Shah et al., 2016; Sarvaiya and Kothari, 2017]. However, whether this trait of sound-responsiveness is widely distributed among microbial word, that remains to be explored; which requires subjecting a larger number of microorganisms to a variety (e.g. in terms of intensity, frequency, duration, etc.) of sonic stimulation, and then studying their response at molecular, cellular, and population level. In such an effort in this study we exposed a gram-negative bacterium namely *Xanthomonas campestris* to a mono-frequency sound of 1000 Hz at different intensities, and the sonic-stimulated bacterial culture exhibiting enhanced exopolysaccharide (EPS) production was subjected to whole transcriptome analysis to know how sonic stimulation has affected bacterial gene expression.

## 2. Materials and Methods

### 2.1. Bacterial strain

*X. campestris* culture was procured from Microbiology Dept., Gujarat University. It was maintained on TYE agar (HiMedia, Mumbai). During sonic stimulation, this bacterium was grown in TYE broth supplemented with 1% w/v dextrose (Merck, Mumbai) and 0.7g/L CaCl_2_ (HiMedia), and incubation was carried out at room temperature.

### 2.2. Sonic stimulation of the bacterial culture

Bacterial culture was subjected to sonic stimulation as described by us previously [Joshi et al., 2018]. The required sound beep of 1000 Hz was generated using NCH^®^ tone generator. Effect of this sound was tested at three different intensities i.e. 40 dB, 66 dB, and 90 dB. The sound file played during the experiment was prepared using Wave Pad Sound Editor Masters Edition v.5.5, in such a way that there is a time gap of one second between two consecutive beep sounds.

Inoculum of the test bacterium was prepared from its activated culture, in sterile normal saline. Optical density (OD) of the inoculum was adjusted to 0.08–0.10 at 625 nm (Agilent Technologies Cary 60 UV-Vis, Bengaluru) to make it equivalent to the McFarland 0.5 turbidity standard.The flasks (Actira;250 mL) containing 100 mL of growth medium (including 5% v/v inoculum) were placed in a wooden chamber. A speaker (Lenovo M0520) was put in this wooden chamber at the distance of 15 cm from the inoculated flasks. Sound delivery from the speaker was provided throughout the period of incubation (72 h). One layer of loose-fill shock absorber polystyrene was filled surrounding the door of the wooden chamber, to minimize any possible leakage of sound from inside the chamber, and also to avoid any interference from external sound. Similar chamber was used to house the ‘control’ (i.e. not subjected to sound stimulation) group flasks. One speaker was also placed in the wooden chamber used for the control flasks at a distance of 15 cm, where no electricity was supplied and no sound was generated [Kothari et al., 2018]. Intensity of the sound was measured with a sound-level meter (ACD Machine Control Ltd, Mumbai) at a distance of 15 cm from the speaker. Background sound intensity was measured to be 38-50 dB.

Intermittent mixing of the contents of the flasks to minimize heterogeneity was achieved by putting the flasks at an interval of every 3 h on a shaker for 15 min (120 rpm). Whenever the flasks were taken out of the wooden chamber, positions of the flasks of a single chamber were interchanged, and their direction with respect to the speaker was changed by 180° rotation. This was done to ensure almost equal sound exposure to all the flasks.

### 2.3. Growth and EPS estimation

At the end of incubation, after quantifying the cell density at 625 nm, the culture was subjected to extraction and quantification of the EPS. For this, culture broth was centrifuged for 10 min at 7500 rpm, and the cell-free supernatant (CFS) was utilized for EPS quantification as per the method described by Li et al. (2012) with some modification. Briefly, 60 mL of chilled acetone was added to 30 mL of CFS, and allowed to stand for 30 min. The EPS precipitated thus was separated by filtration through pre-weighed Whatman # 1 filter paper (Axiva, Haryana). Filter paper was dried at 55°C for 24 h, and weight of EPS on paper was calculated (after ensuring complete evaporation i.e. no further decrease in weight of the filter paper being dried).

## 3. Whole Transcriptome Analysis

Sonic-stimulated *X. campestris* (along with control culture) was subjected to whole transcriptome analysis, so that a holistic picture regarding its response to sonic stimulation can emerge.

### 3.1. RNA isolation, library preparation, and sequencing

RNA was extracted from bacterial pellet (late logarithmic phase) using Hi PURA Bacterial RNA Purification kit (HiMedia, USA) followed by measuring its concentration using QubIT, and QC analysis with RNA 6000 NanoBioanalyzer kit (Agilent, Germany). Whole transcriptome library was prepared from QC-passed samples (RNA integrity number >7) employing NEB next ultra RNA library preparation kit (NEB, USA). Briefly, 10 µg of Total RNA was taken for ribosomal RNA depletion using Ribominus Bacteria kit module (Invitrogen Inc., USA). The rRNAdepleted samples were fragmented through enzymatic method. 1^st^strandcDNA synthesis was achieved using random primers followed by 2^nd^ strand cDNA synthesis, end repair and adapter ligation. The adapter-ligated libraries were multiplexed by adding index sequences via amplification. The adapter ligated and indexed libraries were quantitated by QubIT and validated employing Agilent HS kit. Resultant validated libraries were pooled in equimolar proportion and subjected to sequencing on NextSeq500 platform (Illumina, USA) applying 2×150bp chemistry. Raw data was thenfurther processed after necessary quality check with a mean Q30>70%. All the raw sequence data has been submitted to Sequence Read Archive (SRA). Relevant accessions no. isPRJNA407853 (https://www.ncbi.nlm.nih.gov/bioproject/?term=PRJNA407853).

### 3.2. Genome annotation and functional analysis

Read quality check analysis and mapping: QC report of sequences obtained was produced through FastQC application. High-quality reads were then mapped to the reference genome of *X. campestris* (NCBI accession no. AE008922.1) applying CLC RNASeq analysis protocol of CLC Genomics Workbench version 9.0.

Differential gene expression analysis: The count data was compared between the control and experimental samples to identify the differently expressed genes. The number of reads mapping to each gene was summarized as count data. The count data was first normalized applying quantile normalization tool in CGWB version 9, and then was further statistically analyzed using Kal’s Z-test integrated into CGWB. Genes with p≤0.05were filtered as either up- or down-regulated genes, which were then looked forgene ontology classification in following databases: NCBI gene database (https://www.ncbi.nlm.nih.gov/nuccore/AE008922.1); KEGG (Kyoto Encyclopedia of Genes and Genomes: https://www.genome.jp/dbget-bin/get_linkdb?-t+genes+gn:T00083).

## 4. Results

*X. campestris* was exposed to 1 KHz sound beep at three different intensities, whose effect on its growth and EPS production is presented in Table 1. Since EPS produced by this bacterium is a product of industrial importance and its production was enhanced in *X. campestris* culture subjected to 1000 Hz sound at 66 dB, by 1.69 fold (i.e. 69.30%); we subjected this sonic-stimulated culture to whole transcriptome analysis to have a holistic idea of this bacterium’s response to sound stimuli at molecular level.

**Table 1:** Effect of 1000 Hz sonic stimulus at three different intensities on growth and EPS production by *X. campestris*.

| Sr. no | Sound intensity (dB) | Growth<br>(Cell Density as OD<sub>625</sub>; M±SD)<br>(A) |  |  | EPS (g/L; M±SD)<br>(B) |  |  | EPS Unit (M±SD)<br>(B/A) |  |  |
| --- | --- | --- | --- | --- | --- | --- | --- | --- | --- | --- |
|  |  | C | E | % change | C | E | % change | C | E | % change |
| 1 | 40 | 0.65±<br>0.02 | 0.64±<br>0.01 | -2.53±<br>6.75 | 1.62±<br>0.42 | 0.87±<br>0.37 | -46.19±<br>39.47 | 2.48±<br>0.60 | 1.36±<br>0.55 | -44.79±<br>37.47 |
| 2 | 66 | 0.41±<br>0.01 | 0.46±<br>0.03 | 13.41±<br>4.71 | 2.21±<br>0.18 | 4.23±<br>0.11 | 91.70**±<br>11.05 | 5.39±<br>0.27 | 9.11±<br>0.44 | 69.01**±<br>16.82 |
| 3 | 90 | 0.57±<br>0.00 | 0.67±<br>0.0 | 17.54**±<br>1.43 | 30.53±<br>0.02 | 30.40±<br>0.08 | -0.41±<br>0.19 | 61.95±<br>0.82 | 52.94±<br>0.15 | -14.53**±<br>0.89 |

\*\*p ≤ 0.01; EPS unit was calculated to nullify any effect of change in cell density; C: Control; E: Experimental. All experiments were done in duplicate, and the results presented are mean of two independent experiments. Statistical significance in terms of p-value was assessed through student's t-test done using Microsoft Excel®.

A comparative analysis of ‘control’ *X. campestris* culture with that exposed to sonic stimulation (1000 Hz; 66 dB) in context of gene expression at the whole transcriptome level, resulted in identification of 4,242 genes; of which 961(22.65% of the genome) were expressed differentially in the sonic-stimulated culture at *p*≤0.05. However, to have greater confidence in our interpretation of the data, we applied the dual criteria of *p* ≤ 0.05 and fold-change value ≥1.5. Of the 115 genes following these dual criteria, 57 were down-regulated (Table-2), and 58 were up-regulated (Table-3). Fold change values of the up-regulated genes ranged till11, and those for down-regulated ones ranged till 13. Information on function of all these differentially expressed genes was sourced from KEGG database.A function-wise categorization of significantly differentially expressed genes is presented in Figure 1.

**Table 2:** List of down-regulated genes.

| Sr. No | Feature ID | Fold Change | P-value | Coding for |
| --- | --- | --- | --- | --- |
| 1. | XCC0126 | -13 | 0.02 | dCMP deaminase [EC:3.5.4.12] \| (RefSeq) deoxycytidylate deaminase |
| 2. | XCC2875 | -8.25 | 0.0007 | histone H1 |
| 3. | XCC3989 | -6 | 0.05 | hypothetical protein |
| 4. | XCC3307 | -4 | 0.05 | putative transposase \| (RefSeq) IS1477 transposase |
| 5. | *nuoK* | -3.7 | 3.25E-11 | NADH-quinone oxidoreductase subunit K [EC:1.6.5.3] \| (RefSeq) nuoK; NADH dehydrogenase subunit |
| 6. | *gIII_1* | -3.6 | 0.01 | gIII; adsorption protein |
| 7. | XCC1421 | -2.5 | 0 | hypothetical protein |
| 8. | XCC1181 | -2.2 | 0.0004 | hypothetical protein |
| 9. | XCC3580 | -2.2 | 0.03 | IS1477 transposase |
| 10. | *wxcD* | -2.1 | 4.05E-08 | hypothetical protein |
| 11. | XCC2374 | -2.1 | 0 | hypothetical protein |
| 12. | XCC0930 | -2.0 | 0.02 | hypothetical protein |
| 13. | *vapI* | -2.0 | 1.11E-06 | antitoxin HigA-1 \| (RefSeq) vapI; virulence associated protein |
| 14. | *rpmJ* | -1.9 | 0.01 | large subunit ribosomal protein L36 \| (RefSeq) rpmJ; 50S ribosomal protein L36 |
| 15. | *int_2* | -1.9 | 0.01 | intS; phage-related integrase |
| 16. | XCC4002 | -1.9 | 1.56E-08 | hypothetical protein |
| 17. | XCC2319 | -1.9 | 0.001 | hypothetical protein |
| 18. | XCC1246 | -1.9 | 0.01 | hypothetical protein |
| 19. | XCC1064 | -1.9 | 1.2E-07 | hypothetical protein |
| 20. | XCC3734 | -1.8 | 0.03 | tRNA Met \| (RefSeq) tRNA-Met |
| 21. | XCC4195 | -1.7 | 0.001 | hypothetical protein |
| 22. | *blaI* | -1.7 | 0.03 | BlaI family transcriptional regulator |
| 23. | *smpB* | -1.7 | 0.0008 | SsrA-binding protein |
| 24. | XCC3127 | -1.7 | 0.02 | hypothetical protein |
| 25. | XCC0128 | -1.7 | 0.05 | IS1481 transposase |
| 26. | XCC2091 | -1.7 | 1.38E-05 | tRNA-Leu |
| 27. | *mocA_2* | -1.7 | 0.006 | rhizopine catabolism protein |
| 28. | *rpmD* | -1.7 | 0.005 | 50S ribosomal protein L30 |
| 29. | XCC1300 | -1.7 | 0.0001 | hypothetical protein |
| 30. | *metE* | -1.6 | 1.08E-08 | LysR family transcriptional regulator, regulator for metE and metH \| (RefSeq) metR; MetE/MetH |
| 31. | XCC2019 | -1.6 | 0.01 | hypothetical protein |
| 32. | XCC1636 | -1.6 | 0.05 | IS1480 transposase |
| 33. | XCC1956 | -1.6 | 5.88E-14 | hypothetical protein |
| 34. | XCC1070 | -1.6 | 7.93E-05 | hypothetical protein |
| 35. | XCC0732 | -1.6 | 0.0009 | hypothetical protein |
| 36. | XCC3568 | -1.6 | 0 | hypothetical protein |
| 37. | *exsF_2* | -1.6 | 0.01 | regulatory protein |
| 38. | XCC3979 | -1.6 | 0.0006 | hypothetical protein |
| 39. | *hup* | -1.6 | 0.001 | DNA-binding protein HU-beta \| (RefSeq) hupB; histone-like protein |
| 40. | XCC4080 | -1.6 | 1.03E-11 | polyvinyl alcohol dehydrogenase |
| 41. | XCC2351 | -1.5 | 0 | hemerythrin \| (RefSeq) hypothetical protein |
| 42. | XCC2716 | -1.5 | 2.47E-10 | putative redox protein \| (RefSeq) hypothetical protein |
| 43. | XCC0707 | -1.5 | 0.03 | hypothetical protein |
| 44. | *ubiH* | -1.5 | 1.45E-10 | 2-octaprenyl-6-methoxyphenol hydroxylase [EC:1.14.13.-] \| (RefSeq) ubiH; 2-octaprenyl-6-metho |
| 45. | *rfbD_1* | -1.5 | 0.009 | dTDP-4-dehydrorhamnose 3,5-epimerase [EC:5.1.3.13] \| (RefSeq) rfbD; strX protein |
| 46. | XCC0309 | -1.5 | 0.009 | hypothetical protein |
| 47. | *trx_1* | -1.5 | 0.005 | thioredoxin reductase (NADPH) [EC:1.8.1.9] \| (RefSeq) trxB; thioredoxin reductase |
| 48. | XCC0099 | -1.5 | 0.003 | hypothetical protein |
| 49. | XCC3560 | -1.5 | 0.03 | hypothetical protein |
| 50. | XCC1587 | -1.5 | 0.02 | methionine sulfoxide reductase heme-binding subunit \| (RefSeq) sulfite oxidase subunit YedZ |
| 51. | XCC0242 | -1.5 | 0.007 | hypothetical protein |
| 52. | XCC4047 | -1.5 | 1.45E-07 | hypothetical protein |
| 53. | *rhsD_2* | -1.5 | 0.0008 | RhsD protein |
| 54. | *pqqC* | -1.5 | 2.59E-05 | pyrroloquinoline-quinone synthase [EC:1.3.3.11] \| (RefSeq) pqqC; pyrroloquinoline quinone bio |
| 55. | *rpmF* | -1.5 | 0.03 | large subunit ribosomal protein L32 \| (RefSeq) rpmF; 50S ribosomal protein L32 |
| 56. | XCC2668 | -1.5 | 0.0007 | hypothetical protein |
| 57. | XCC0709 | -1.5 | 0.001 | inner membrane protein \| (RefSeq) hypothetical protein |

**Table 3:** List of up-regulated genes.

| Sr. No. | Feature ID | Fold Change | P-value | Coding for |
| --- | --- | --- | --- | --- |
| 1. | XCC3137 | 11 | 0.04 | Hypothetical protein |
| 2. | XCC1609 | 7.5 | 0.02 | Transposase, IS5 family |
| 3. | XCC1102 | 4.4 | 0.001 | Hypothetical protein |
| 4. | XCC0326 | 3.6 | 4.89E-05 | Hypothetical protein |
| 5. | XCC3640 | 3.2 | 0.01 | Hypothetical protein |
| 6. | XCC1628 | 2.6 | 0.0004 | IS1477 transposase |
| 7. | XCC2591 | 2.6 | 0.03 | Hypothetical protein |
| 8. | XCC3924 | 2.5 | 0.04 | Hypothetical protein |
| 9. | XCC3934 | 2.5 | 0.0004 | Hypothetical protein |
| 10. | XCC1065 | 2.5 | 0.007 | Hypothetical protein |
| 11. | *exsF_1* | 2.2 | 0.003 | two-component system regulatory protein |
| 12. | XCC3193 | 2.2 | 1.4E-06 | Hypothetical protein |
| 13. | XCC0461 | 2.1 | 0.002 | Hypothetical protein |
| 14. | *fimT_2* | 2 | 0.04 | Type IV fimbrial biogenesis protein |
| 15. | XCC0102 | 1.9 | 0.003 | Hypothetical protein |
| 16. | XCC0136 | 1.9 | 0.01 | IS1480 transposase |
| 17. | XCC1463 | 1.8 | 0.002 | Hypothetical protein |
| 18. | XCC2104 | 1.8 | 0.0007 | Hypothetical protein |
| 19. | *trbP* | 1.8 | 0.03 | Conjugal transfer protein |
| 20. | XCC2117 | 1.8 | 1.6E-06 | Protein-tyrosine phosphatase (low molecular weight phosphotyrosine ) |
| 21. | XCC2979 | 1.8 | 7.31E-08 | Hypothetical protein |
| 22. | XCC1798 | 1.7 | 0.02 | IS1481 transposase |
| 23. | XCC3549 | 1.7 | 0.01 | Hypothetical protein |
| 24. | *pnuC* | 1.7 | 1.24E-10 | Nicotinamide mononucleotide transporter |
| 25. | *sdhC* | 1.7 | 0.008 | Succinate dehydrogenase / fumarate reductase, cytochrome b subunit |
| 26. | XCC2957 | 1.7 | 5.16E-07 | Hypothetical protein |
| 27. | XCC0540 | 1.6 | 0.002 | Hypothetical protein |
| 28. | XCC1687 | 1.6 | 0.0003 | General stress protein |
| 29. | *cobD* | 1.6 | 7.28E-10 | Adenosylcobinamide-phosphate synthase |
| 30. | *tufA* | 1.6 | 0 | Elongation factor Tu) |
| 31. | XCC1784 | 1.6 | 5.77E-06 | Transporter |
| 32. | *gIII_2* | 1.6 | 0.0007 | Adsorption protein |
| 33. | XCC0588 | 1.6 | 0.008 | Hypothetical protein |
| 34. | XCC1669 | 1.6 | 0.0005 | Cytochrome b561 |
| 35. | *rubA* | 1.6 | 0.03 | Rubredoxin |
| 36. | XCC0840 | 1.6 | 3.56E-09 | Hypothetical protein |
| 37. | XCC3266 | 1.6 | 0.0003 | Hypothetical protein |
| 38. | *acvB_2* | 1.6 | 1.37E-11 | virulence protein |
| 39. | *btuE_1* | 1.6 | 2.85E-05 | Glutathione peroxidase; ABC transporter vitamin B12 uptake permease |
| 40. | XCC2561 | 1.6 | 2.58E-05 | Hypothetical protein |
| 41. | XCC3799 | 1.6 | 0.005 | Hypothetical protein |
| 42. | *mcrB* | 1.5 | 7.27E-06 | 5-methylcytosine-specific restriction enzyme B |
| 43. | *mmsB* | 1.5 | 9.76E-11 | 3-hydroxyisobutyrate dehydrogenase |
| 44. | *virB1* | 1.5 | 0.01 | Type IV secretion system protein |
| 45. | *rbfA* | 1.5 | 0.001 | Ribosome-binding factor A |
| 46. | *hutU* | 1.5 | 0 | Urocanatehydratase |
| 47. | XCC3085 | 1.5 | 0.02 | Hypothetical protein |
| 48. | XCC0281 | 1.5 | 0.008 | Hypothetical protein |
| 49. | XCC0717 | 1.5 | 0.008 | Cell division protein MraZ |
| 50. | *mexA* | 1.5 | 0 | Membrane fusion protein, multidrug efflux system |
| 51. | XCC3712 | 1.5 | 0.01 | Hypothetical protein |
| 52. | XCC3795 | 1.5 | 0.005 | Hypothetical protein |
| 53. | XCC2967 | 1.5 | 0.0004 | Site-specific DNA-methyltransferase (adenine-specific) |
| 54. | *exsG* | 1.5 | 0.02 | Two-component system sensor protein |
| 55. | *murG* | 1.5 | 0 | Membrane-associated glycosyltransferase involved in peptidoglycan biosynthesis |
| 56. | XCC4142 | 1.5 | 0.003 | Type I restriction enzyme M protein [EC:2.1.1.72] \| (RefSeq)<br>XmnImethyltransferase |
| 57. | *rpsT* | 1.5 | 0.007 | Small subunit ribosomal protein S20 |
| 58. | *fdsC* | 1.5 | 0.003 | Formate dehydrogenase accessory protein |

**Figure 1:**
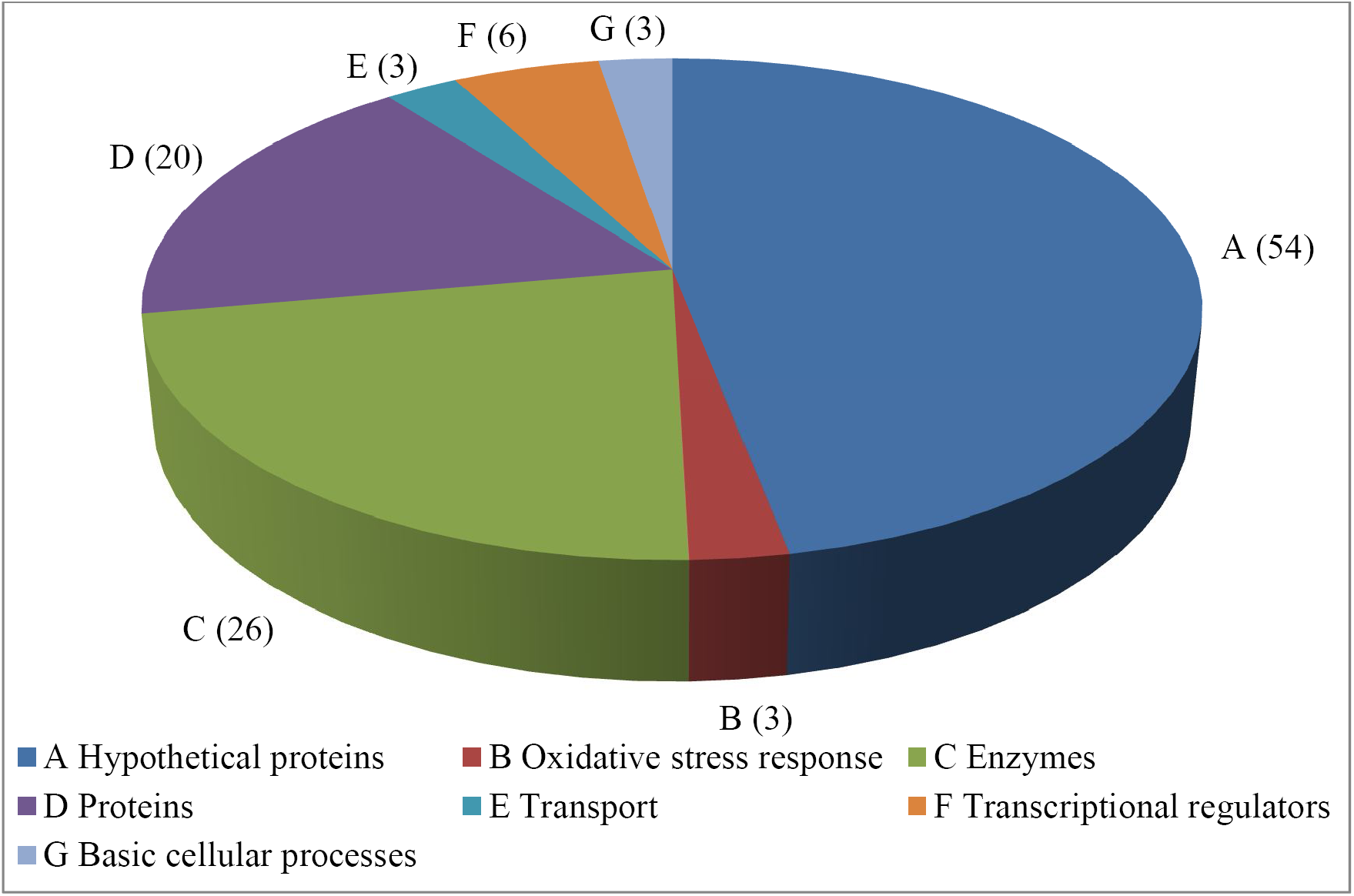
Function-wise categorization of the differentially expressed genes.

Of the 115 differentially expressed genes (DEG), a total of 54 were hypothetical proteins with no predicted functions. Possibly quite a few of them may be the genes involved in bacterial response to vibrations/ sonic stimulation, and microbial response to vibrational/ sonic/ mechanical stimulation been an under-investigated area, these genes have not been assigned any particular function yet. Excluding the hypothetical proteins, there were a total of 61 genes with known functions in the differentially expressed (fold change ≥1.5) category. Since the sonic-stimulated culture overproduced EPS, we tried to see whether the genes involved in xanthan biosynthesis and/or excretion are getting expressed differently. Genes involved in xanthan biosynthesis are part of the *gum* operon. Though none of the gum genes got differently expressed in the sonic-stimulated culture satisfying the cut-off of 1.5 fold; five of them (*gumC* 1.27 fold ↓, *gumG* 1.36 fold ↓, *gumP* 1.48 fold↓, *gumH* 1.16 fold ↑, *gumI* 1.25 fold ↑) got differently expressed at fold change values ranging from 1.16-1.48 fold. Of them, *gumH* is responsible for addition of the internal α-1, 3-mannose, and *gumI* for addition of the terminal β-1, 4-mannose [Becker et al., 1998]. *gumC* is among the genes whose inactivation in a wild-type background is believed to be lethal. Its product may be needed for polymerization orexport of the polymer [Katzen et al., 1998]. Synthesis of many extracellular enzymes and xanthan is reported to be activated by the products of the rpf (regulation of pathogenicity factors) gene cluster [da Silva et al., 2002]. In our sonic-stimulated culture, a total of three rpf genes (*rpf B, rpfG, rpfI*) were differently expressed, albeit at fold change values <1.5. Inactivation of rpfI is believed to reduce expression levels of proteases and endoglucanases [Lee et al., 2005]. Another EPS-associated gene getting expressed differently was *rfbD* (1.5 fold↓). *RfbD* mutants of *X. campestris* may accumulate an unusually high level of dTDP-glucose, causing competitive inhibition of UDP-glucose pyrophosphorylase, resulting in lower levels of UDPglucoseand, hence, reduced EPS production [Köplin et al., 1993]. Among the upregulated genes, we could find two such genes (XCC2117 1.8 fold↑, *murG* 1.5 fold ↑) which have been shown to be contributing towards bacterial polysaccharide synthesis. Of them, XCC2117 codes for protein-tyrosine phosphatase, and tyrosine phosphorylation is recognized as a key regulator of bacterial physiology [Standish and Morona, 2014].

Focusing exclusively on the genes expressed differentially with fold-change values ≥1.5, we found two genes (*exsF* and *exsG*) belonging to the two-component system getting upregulated. Two-component systems are involved in transduction of extracellular signal [Wojnowska et al., 2013] (e.g. external sonic stimulation, in case of current study), and responding to it. A total of 8 transposases were also expressed differently in the experimental (i.e. sonic stimulated) culture. Organisms harboring ISs (Insertion Sequences) are subject to a variety of mechanisms that enhance genomic plasticity [Cesbron et al., 2015]. It may be speculated that in face of the mechanical stress created by sonic vibrations, the bacterium might have responded by modulating its genomic plasticity. Two genes (*virB1; mexA*) involved in secretion/ efflux were also upregulated, which might have contributed to enhanced export of the EPS by the experimental culture.

Six transcriptional regulators experienced altered expression owing to sonic stimulation, which in turn can be expected to have affected expression of multiple genes/ traits regulated by them. One such regulator of the LysR family-*metE* was downregulated by 1.6 fold. XCC3734 coding for tRNA Met was down regulated by 1.8 fold. Methionine is known to be required for the correct functioning of diverse cellular processes such as translation of mRNA into protein [Liu et al., 2013]. XCC1587 coding for methionine sulfoxidereductaseheme-binding subunit was also downregulated (1.5 fold↓). This might have resulted in a reduction in bacterium’s (who can be said to be facing the challenge of mechanical stress) ability to withstand oxidative stress, and tomaintainenvelopeintegrity[https://string-db.org/newstring_cgi/show_network_section.pl?identifier=YedZ].

If we compare the effect of sonic stimulation on the gram-negative bacterium *X. campestris* described in this study at the whole transcriptome level, with our earlier such study [Joshi et al., 2018] with another gram-negative bacterium *Chromobacterium violaceum*, the major categories of genes getting differently expressed in both cases include those coding for transcriptional regulators, enzymes, transposases, and a variety of proteins. Though we could not draw any major generalized conclusions, from the differential gene expression profile of the sonic-stimulated *X. campestris* culture described in this study, which can explain the molecular basis associated with enhanced EPS production of this bacterium, or the mechanism of how it perceives and responds to the sonic vibrations; this report along with our previous reports [Kothari et al., 2014; Sarvaiya and Kothari, 2015; Kothari et al., 2016; Sarvaiya and Kothari, 2017; Joshi et al., 2018] in this field do provide further indication of the property of ‘responsiveness to sonic-stimulation’ being present in multiple microorganisms (bacteria and yeast).

## Acknowledgement

Authors thank Nirma Education and Research Foundation (NERF) for infrastructural support; GUJCOST (Gujarat Council on Science and Technology) for financial support; Vidhi Shah and Jinal Sukhadiya for assistance in lab; and Microbiology Department, Gujarat University for providing the bacterial culture.

